# PIN7 auxin carrier is a terminator of radial root expansion in *Arabidopsis thaliana*

**DOI:** 10.1101/253807

**Authors:** Michel Ruiz Rosquete, Jürgen Kleine-Vehn

## Abstract

Directional growth of lateral roots is critical for radial expansion and soil coverage. Despite its importance, almost nothing is known about its molecular determinants. Initially, young lateral roots (LRs) grow away from the parental root maintaining the angle acquired shortly after emergence. A second downwards bending response to gravity terminates the so-called plateau phase and thereby limits the radial root expansion. Here we show that the exit from the plateau phase correlates with an increase in auxin signalling at the tip of LRs. Moreover, the increase in auxin levels induces the termination of the plateau phase, which requires PIN auxin efflux carriers. Our data suggests that the developmental increase of auxin triggers the preferential de-repression of PIN7 in gravity-sensing columella cells. The subsequent polarization of PIN7 heralds the bending towards gravity and, hence, the exit from the plateau phase. This developmental framework reveals the distinct roles of PIN auxin efflux carriers in controlling the radial growth of root systems.

## Introduction

Secondary roots comprise the major biomass of mature root systems and, hence, their growth and orientation exert a large influence on the overall plant performance. Lateral roots (LR) differ from primary roots in their orientation with respect to the gravity vector, captured by the gravitropic set point angle (GSA) [1]. Initially, LRs grow away from the main axis of the primary root at non-vertical orientations, thus facilitating accessibility to the soil for water and nutrient acquisition and increasing mechanical support [2].

In roots, gravity is perceived in the columella cells of the root cap but the actual growth response to gravity is achieved in the elongation zone. The growth regulator auxin provides the mobile link between the two parts [3–6]. This is achieved, in large part, thanks to the activity and polarized expression of several auxin transporters of the PIN family. In the main root, polar/asymmetric localization of PIN3, PIN4 and PIN7 in columella cells aligns the auxin transport towards gravity, which ultimately leads to increased auxin at the lower root flank, inducing differential root growth and bending towards gravity [6].

Young, stage I lateral roots emerge from the main root at a 90°-degree angle, subsequently responding to gravity in stage II. This downwards bending is highly controlled, establishing an initial gravitropic set-point angle (GSA) of approximately 62° (Fig 1A) [7]. In this early phase of lateral root development, only PIN3 is expressed in the LR columella cells, where it establishes a relatively weak asymmetric auxin signalling [7; 8], which may in part explain the partial response to gravity of LRs [7]. After this first gravitropic response, PIN3 expression diminishes and LRs continue to grow maintaining the acquired GSA [7]. We previously coined the term “plateau phase” for developmental stage III of LRs [7].

**Figure 1.**
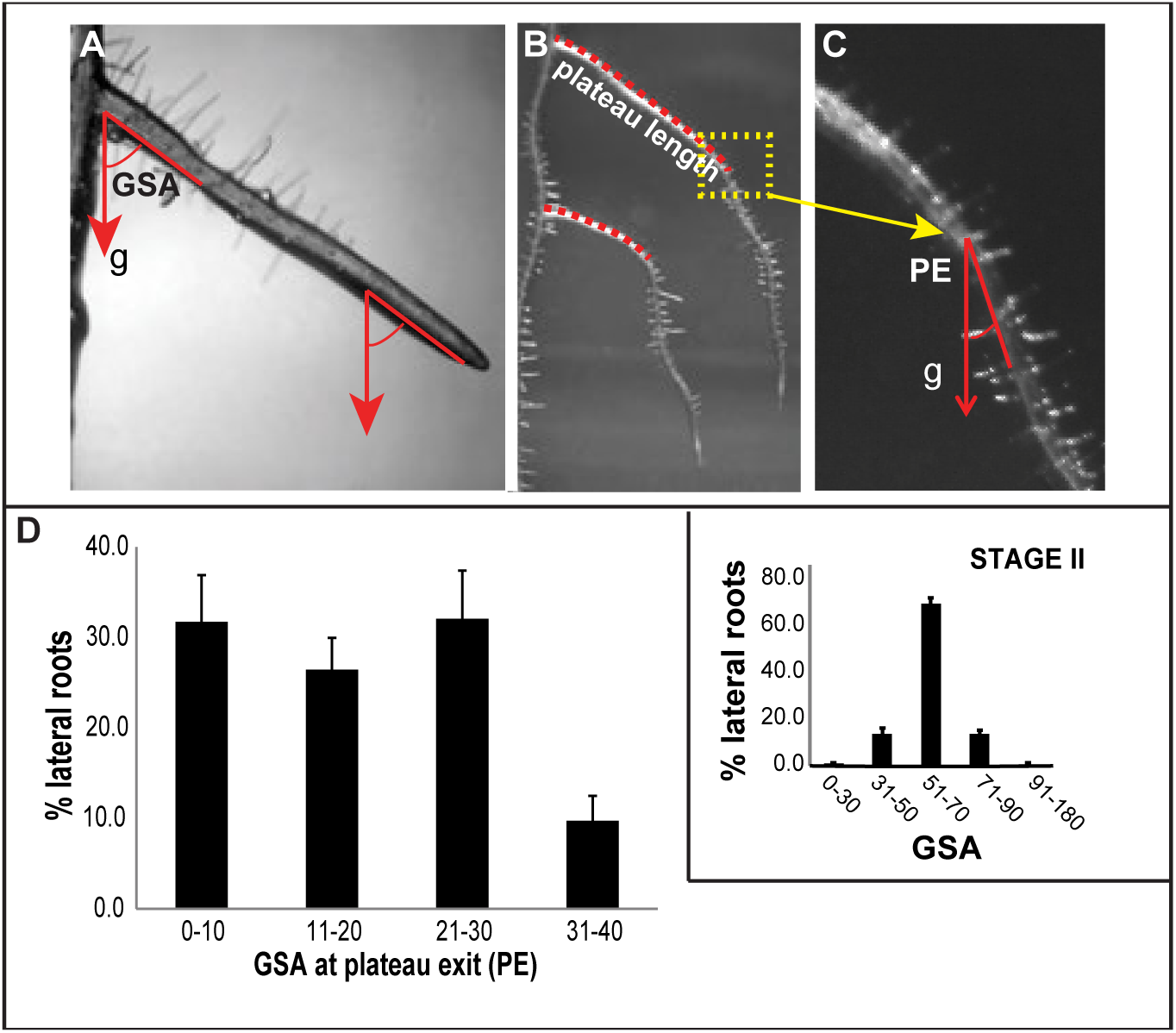
Gravitropic response at plateau exit is highly variable. A) During the plateau phase or stage III, lateral roots (LRs) grow further from the parental root while keeping the initial gravitropic set-point angle (GSA) acquired in stage II. Arrows denote the gravity (g) vector. B) Two representative stage IV LRs are shown. Discontinuous red lines mark the length of their plateau trajectories. C) Enlargement of the transit from stage III to IV, for the upper LR in B. D) PE distribution of Col-0 (wild type) LRs. Error bars represent SEM (n=6 experiments; each experiment with n≥20 seedlings (40–60 LRs)). Distribution of stage II GSA values for Col-0 [see 7] is provided in the inset.

The initial establishment of the GSA and the length of the plateau phase determine the radial expansion of the root system. It currently remains unknown how this plateau-like growth phase of LRs is terminated.

## Results

### Plateau length of lateral roots contributes to root system architecture

The exit from the plateau phase is characterized by a second, positive gravitropic response, lowering the GSA to a value typically below 30˚ in stage IV LRs (Fig 1B-D). The GSA at the plateau exit (PE) fluctuated within a broad range (0°-30°) (Fig 1C, D), suggesting that this developmental response is highly variable compared to the establishment of the initial GSA at stage II [7] (inset in Fig 1D). The initial GSA, the length of the plateau phase and its corresponding PE angle jointly determine the radial expansion of the root system.

Given the importance of the plateau length for root system architecture, we focus here on its regulation. Most individual LRs displayed a clearly distinguishable plateau phase; however, we also observed individual laterals that did not display a clear plateau but rather showed a continuous decline of GSA throughout development (Fig 2A, upper LR). Although their frequency was rather low (5-10%) (Fig 2E), it needs to be seen whether this phenomenon reflects an alternative developmental program or is due to some substrate constraint. On the other hand, wavy growth without major changes in overall growth direction (GSA) is often observed during the plateau phase (Fig 2B, C). Waviness of *Arabidopsis* roots has been suggested to result from the mechanical torsion of the root at the 2D agar surface [9]. In agreement, we found that the frequency of “wavy” lateral roots diminishes by approximately 40% when seedlings were grown within the gel of cylindrical vessels (3D *in vitro* system) (Fig 2D, E).

**Figure 2.**
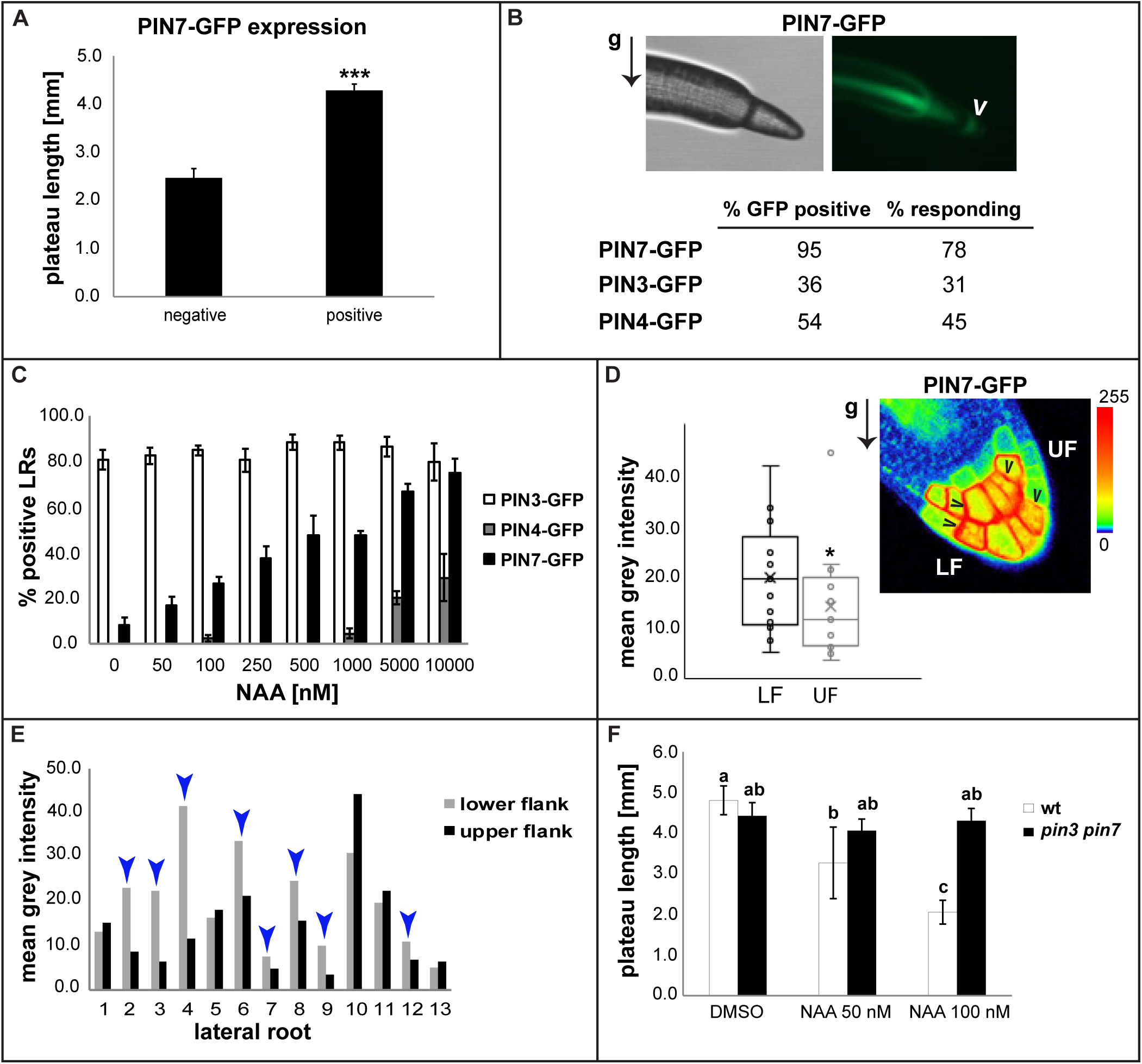
Characterization of the plateau phase. A) A low percentage of LRs display a continuous decline of GSA throughout development (“no plateau”); notice the contrast between upper LR (no plateau) and bottom LR (LR in frank plateau phase). B, C) Growth during the plateau phase may show some degree of waviness. D) Representative 20-DAG wild type seedling (Col-0), grown in a cylindrical vessel (3D model). E) Incidence of waviness during plateau is lower in seedlings that grow within the agar (cylindrical vessels), compared to seedlings grown on the surface of medium (plates). Percentages from a representative experiment are given (≥150 LRs; ≥30 seedlings). F, G) Plateau segments (red lines in F) of higher order LRs (1-3) are typically longer than those of younger or lower order LRs (4-6 in the example). Average values of plateau length were obtained for higher (1-3) and lower order LRs (4-), for individual Col-0 seedlings (G). Representative experiments are shown. Error bars represent SEM (n>30 seedlings per experiment). Three asterisks denote p<0.001 (Student‘s t-test). Representative 14-DAG Col-0 seedling is shown in F. H) Higher order LRs are longer also in the 3D growth model. A magnification of the seedling in D is shown. Red and yellow vertical lines help to compare the plateau length of a higher order LR (red arrowhead) with that of a younger LR (yellow arrowhead).

When observing the lateral roots along the main roots, we also noticed that positional effects may determine the individual growth behaviour. The developmentally oldest LRs (more proximal to the hypocotyl) had longer plateau segments compared to those that emerge later on during root development (Fig 2F). Under our growth conditions, plateau of the first three uppermost located laterals typically extend 4-5 mm and were substantially longer compared to more distant, younger LRs (Fig 2F, G). Notably, we observed a similar effect in the 3D *in vitro* system (Fig 2H). We concluded that the first emerging LRs are most critical for the overall radial expansion of the root system in *Arabidopsis* (Fig 2F-H). Accordingly, we specifically investigated the plateau length of the first three LRs, which we refer to as “higher order lateral roots”.

### Auxin restricts radial expansion of lateral roots

Lateral roots display an age-dependent intensification of auxin signalling at their tips[10]. However, a possible correlation between increases of auxin signalling and transitions between LR developmental stages has not been explored. To test that possibility, we used the DR5::GFP reporter line to quantify the levels of auxin signalling in the columella region of LRs in all four developmental stages. DR5 activity was significantly augmented from stage I to II, correlating with the onset of auxin-dependent bending towards gravity (Fig 3A, B) [7]. Moreover, we observed a second significant increase in nuclear auxin signalling, which accompanied the transition from stage III to stage IV (Fig 3A, B).

**Figure 3.**
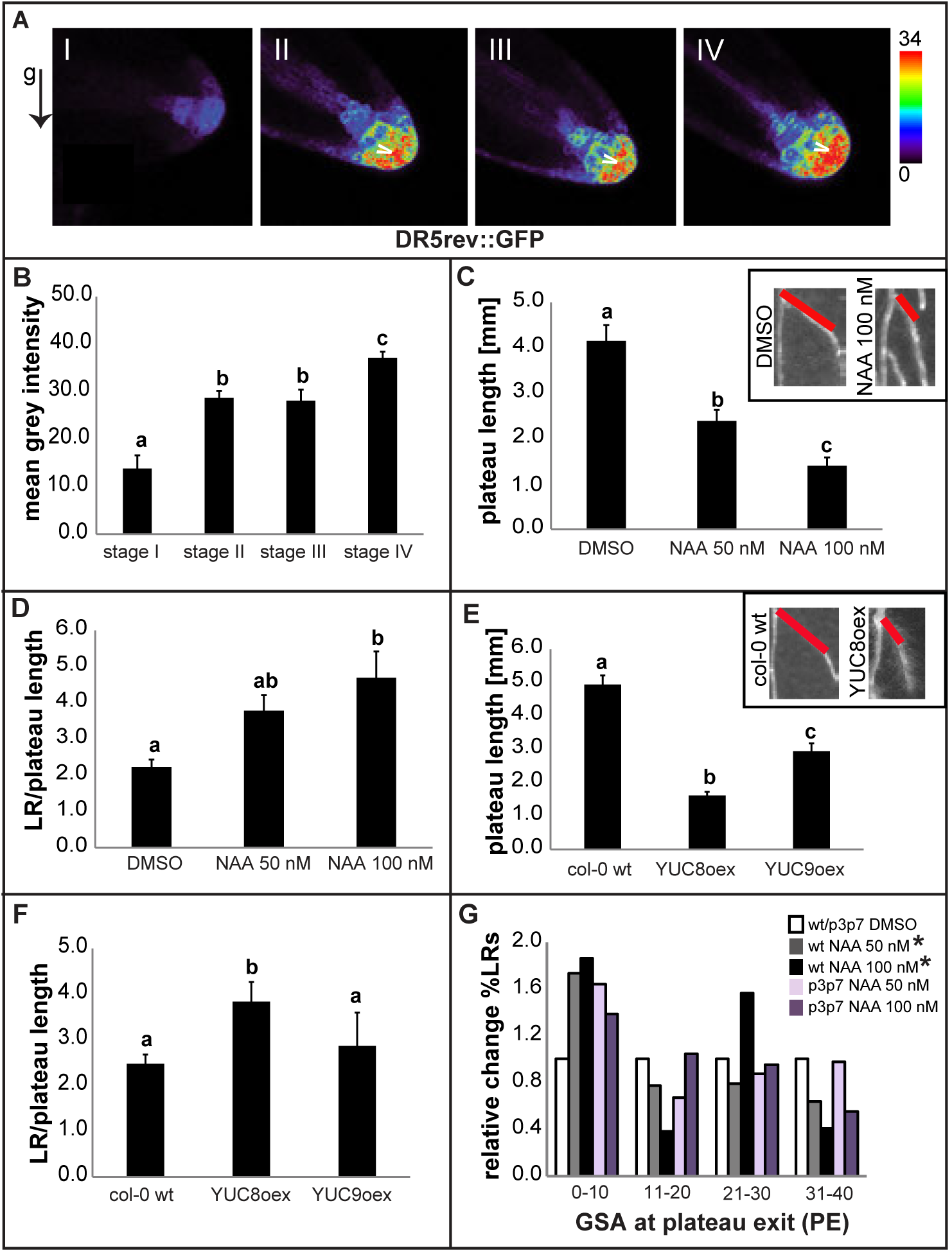
Auxin induces the termination of plateau phase. A, B) Increases of auxin signaling at the root tip, visualized by DR5-driven GFP signal (A), accompany the transitions from stage I to II and from stage III to IV during LR development. Arrowheads denote areas of high DR5 activity. A semiquantitative color-coded heat-map of DR5-GFP signal is provided. An arrow denotes the gravity (g) vector. B) Mean intensity of the DR5-GFP reporter signal was measured in the quiescent center + columella region, in confocal images. A representative experiment is shown. Error bars represent SEM (n≥10 LRs per stage, per experiment). C, D) Treatments of 4-DAG Col-0 wild type seedlings with 50 nM and 100 nM auxin significantly reduce plateau length (C) and increase the LR length/plateau length ratio (D). E, F) Lines overexpressing the auxin biosynthetic genes YUC8 and YUC9 (YUCoex) phenocopy NAA treatments. Insets in C and E illustrate how exogenous (C) and endogenous (E) increases of auxin determine a reduction of plateau length and a steeper plateau exit angle. A representative experiment is shown for both NAA treatments and YUCoex phenotyping. Error bars represent SEM (n>30 LRs per experiment). Statistical significance in A-F was evaluated by one-way ANOVA followed by multiple comparisons Tukey test. Distinct letters indicate statistically significant differences (p<0.05). G) Auxin treatments determine a stronger graviresponse in LRs of wt seedlings, evidenced by the significantly higher percentage of LRs exiting plateau at angles in the lowest category (0-10) of the PE distribution. pin3 pin7 double mutants show slight resistance to the auxin determined shift of the PE distribution. The effect of auxin treatment on the PE distributions of both wt and mutants was expressed as relative change in the percentage of PE values (in each PE category) caused by the treatment, with respect to the percentage of the non-treated controls in the same category. A representative experiment is shown (n>30 LRs per experiment). Error bars represent SEM. One asterisk denotes p<0.05 (Kolmogorov-Smirnov test).

Based on these observations, we assumed that an increase in auxin signalling could play a role in regulating also the second bending response to gravity, which terminates the plateau phase. To gain evidence, we exogenously applied the synthetic auxin NAA in the nanomolar range, which strongly reduced the length of the plateau (Fig 3C). Overall lateral root growth was also reduced in those experiments; however, the reduction of the plateau length was comparatively greater (Figure 3D), suggesting that auxin negatively impacts the radial expansion of root systems by terminating the plateau phase. In agreement, the induction of YUC-dependent auxin biosynthesis largely phenocopied the effect of auxin application, similarly reducing the plateau length of LRs (Fig 3E, F). Notably, auxin also caused a considerably increased frequency of laterals exiting the plateau at steeper angles (between 0˚ and 10˚) (Fig 3G), reflecting a stronger response to gravity. This data suggests that both exogenous application and endogenous increase of auxin levels negatively limit the plateau length of LRs by stimulating their response to gravity.

### Auxin preferentially triggers PIN7 expression in higher order lateral roots

Next, we assessed by which mechanism auxin may affect the gravity responsiveness of plateau lateral roots. Expression of PIN auxin carriers in the columella cells of the root tip is crucial for the root gravitropic response [6]. One of the most intriguing characteristics of the plateau phase is the very low expression levels or even absence of all three columella PINs (PIN3, PIN4 and PIN7) [7]. To assess whether the derepression of PINs heralds the exit from plateau phase, we examined pPIN3:PIN3-GFP, pPIN4:PIN4-GFP and pPIN7:PIN7-GFP expression in higher order lateral roots. Even though expression of the three PINs remained low during the plateau phase, we observed a tendency towards preferential PIN7-GFP de-repression in older and hence longer (≥ 4 mm) stage III lateral roots (Fig 4A). This prompted us to further explore the correlation between columella PIN expression and its requirement to exit from the plateau phase. Therefore, we inspected higher order LRs presumably facing the termination of their plateau phase (≥ 4 mm) (Fig 4B). PIN7 expression was frequent (95%) at the tip of those laterals, whereas expression of PIN3 and PIN4 occurred at much lower frequencies (36% and 54%, respectively). Auxin is known to feedback on PIN expression [11] and auxin signalling increased during lateral root maturation (Fig 3A, B). Therefore, we wondered whether auxin may impact distinctly on PIN7 expression in LRs. To assess this, we performed NAA treatments on stage I and stage II LRs, where PIN4 and PIN7 are strongly repressed [7]. The percentage of laterals expressing PIN4-GFP started to increase only at a very high concentration (1 μM), whereas PIN7-GFP showed steady induction starting at relatively low concentrations (50 nM) (Fig 4C). Hence, we conclude that PIN7 expression in LR columella cells is particularly sensitive to auxin.

**Figure 4.**
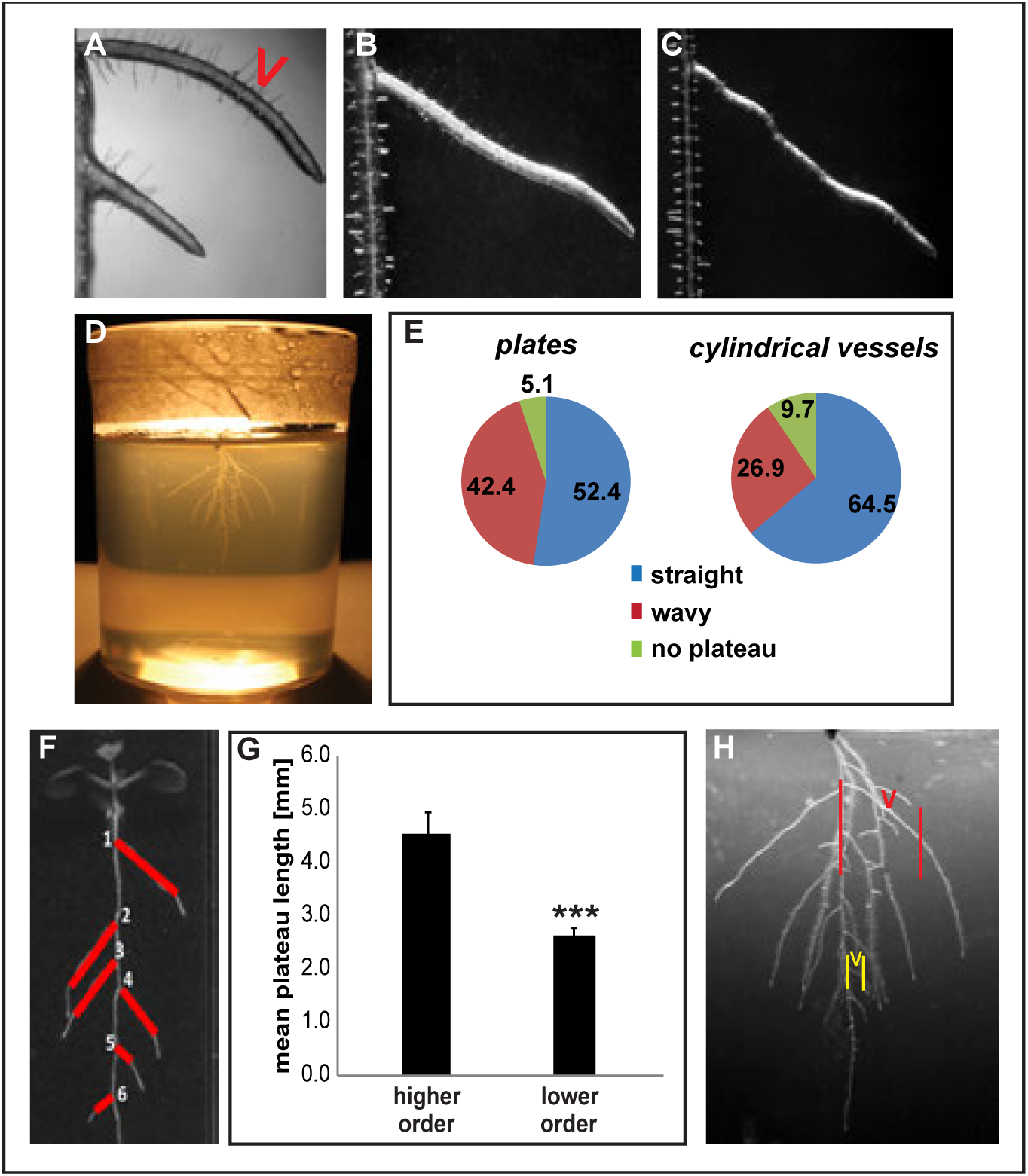
PIN7 has a preferential role in plateau phase termination. A) Examination of higher order LRs (6-8 DAG) revealed a positive correlation between likelihood of PIN7 expression and length of the plateau phase. A representative experiment is shown (n≥60 LRs; n≥30 seedlings per experiment). Three asterisks denote p<0.001 (Student‘s t-test). B) Compared with PIN3 and PIN4, PIN7 is more frequently detected (95%) in advanced stage III LRs (>4 mm). In 78% of tracked cases, expression of PIN7-GFP at that developmental point correlated with the subsequent graviresponse (within 12 hours), leading to exit from plateau. An arrowhead indicates pPIN7-GFP expression at the root tip. An arrow denotes the gravity (g) vector. C) 4-DAG seedlings of PIN3::PIN3-GFP, pPIN4::PIN4-GFP and pPIN7::PIN7-GFP reporter lines were treated for three days with the indicated concentrations of NAA and the presence of detectable signal of the corresponding PIN in the tip of stage I and II LRs was examined. An average of three experiments is shown (n≥30 LRs per experiment). D, E) Expression of PIN7-GFP in columella cells of the root tip exhibits polarization in a high percentage of cases of LRs apparently exiting plateau. Polarization was assessed by measurement of PIN7-GFP fluorescence intensity at lateral plasma membrane domains of the outermost columella cells at the lower flanks (LF) region compared with those at the upper flanks (UF). Arrowheads indicate asymmetric PIN7-GFP distribution. A semiquantitative color-coded heat-map of PIN7-GFP signal is provided (D). Results from a representative experiment are presented (n>10 LRs, n>7 seedlings per experiment) as D) an average of all LRs analyzed and E) mean intensities for UF and LF for all LRs assessed in the experiment, individually. Blue arrowheads in E indicate cases of polarization towards the gravity vector (LF>UF). One asterisk in D denotes p<0.05 (Student‘s t-test). F) pin3 pin7 mutants show resistance to the reduction of the plateau length caused by NAA treatments on the wild type (4-DAG seedlings). A representative experiment is shown (n>30 LRs per experiment). Error bars represent SEM. Statistical significance was evaluated by two-way ANOVA followed by multiple comparisons Tukey test. Distinct letters indicate statistically significant differences (p<0.05).

### Asymmetric PIN7-GFP expression correlates with differential growth process

We next assessed whether PIN de-repression in columella cells could indeed mediate stage transition and, hence, alterations in growth direction of lateral roots. We accordingly tracked individual laterals with PIN-GFP expression, using a fluorescence binocular microscope, and recorded whether they actually altered their growth direction towards gravity within 12 hours (Fig 4B). pPIN3:PIN3-GFP and pPIN4:PIN4-GFP positive stage III LRs displayed a plateau exit in 31% and 45% of cases, respectively. This data suggests that *PIN3* or *PIN4* expression in columella cells is not *per se* initiating gravitropic bending in late stage III LRs. In contrast, pPIN7:PIN7-GFP expression showed a rather high correlation (78%) with alterations in growth direction. This corroborates the central role of PIN7 during termination of the plateau phase, but also raises the questions why some PIN7-GFP positive lateral roots do not display differential growth response.

As PIN7-GFP was most readily expressed in higher order lateral roots and showed the highest correlation with alterations in growth behaviour, we subsequently analysed its subcellular localization, using confocal microscopy. We detected polarization of PIN7-GFP expression in only about 67% of the PIN7-GFP expressing LRs (Fig 4D, E). This is very much in range with the roughly 78% of PIN7-GFP positive lateral roots exiting the plateau phase. Accordingly, we conclude that not PIN7 expression as such, but rather its asymmetric pattern largely correlates with differential growth in LR transiting from stage III to IV. This finding also suggests that auxin-dependent de-repression of mainly PIN7 terminates the radial expansion of the root system.

We observed an endogenous increase of auxin signalling at the transition from stage III to stage IV. Moreover, PIN7 expression shows relatively high auxin sensitivity. Accordingly, we hypothesize that auxin could be the actual developmental trigger to preferentially control PIN7 (and to a lesser extent PIN3 and PIN4) derepression in stage III higher order LRs. This auxin-dependent regulation presumably initiates a PIN-dependent stage transition, reducing radial exploitation of the root system. In line with this, *pin3 pin7* double mutants were less sensitive to the auxin-induced reduction of the plateau length and the associated shift in PEA (Fig 4F and 3G). We therefore conclude that developmentally defined auxin levels determine columella PIN expression in lateral roots, which is, in conjunction with a polarization mechanism, important for the auxin-dependent termination of radial lateral root growth.

## Discussion

Three decades ago, Alistair Fitter noted: “*many roots show a change from a diageotropic (horizontal with respect to the gravity vector) to a positively geotropic response after a period of growth, which results in an increase in the volume of soil exploited*” [12]. By “after a period of growth”, Fitter most likely referred to what we call the plateau phase, an LR developmental stage of relevance for the initial spread of the root system. The molecular mechanisms and players that control the spatial expansion of root systems remain largely unexplored. With the present work, we shed light on how the non-gravitropic growth during the plateau phase is terminated. We show that auxin plays a central role in the restriction of the plateau phase.

For auxin to steer root gravitropic responses, the activity and polar localization of auxin transporters at the plasma membrane are necessary [3–8]. In a previous work, we demonstrated a role for the auxin effluxer PIN3 in the initial gravitropic response of young lateral roots [7]. Our new results propose an extended role for auxin as a negative regulator of the radial expansion of the root system. PIN-dependent auxin transport is similarly required to implement the second response to gravity. The data presented suggest that a different member of the PIN protein family, PIN7, mainly mediates the gravitropic response that determines exit from the plateau phase. Notably, a developmentally defined increase in auxin signalling in the tip of LRs correlates with the transition from stage III to stage IV. We found that exogenously applied auxin preferentially induces the de-repression of PIN7 in stage I and II LRs. Based on this data, it is very likely that the observed rise in auxin signalling preferentially triggers PIN7 expression. In agreement, PIN7 is, compared to PIN3 and PIN4, more readily expressed at the tip of late stage III LRs. Moreover, PIN7 expression and subsequent polarization correlate most strongly with gravitropic response. This set of data proposes a central role of PIN7 in determining LR exit from the plateau phase. Currently, little is known about transcription factors that may differentially impact on columella PIN expression in LRs. Notably, transcription factors MYB88 and FOUR LIPS were suggested to regulate both PIN3 and PIN7 expression in LRs [13]. It accordingly remains to be seen how the preferential and developmentally defined expression of PIN7 is molecularly achieved.

On the other hand, our data also shows that PIN3, PIN4, or PIN7 expression is not sufficient to induce differential growth and that additional polarity cues are required. AGCVIII kinases of the D6 and PINOID families impact on PIN phosphorylation and are known to modulate the localisation and/or transport activity of PINs at the plasma membrane [14–17]. However, very little is known about phosphorylation of PIN7 and its potential contribution to LR growth. Intriguingly. *LAZY/DEEPER ROOTING (DRO)* genes were recently found to regulate columella PIN polarization towards gravity in main and LRs [18–20]. Intriguingly, *DRO1* and *DRO2* show variable, age-dependent expression in LRs of Arabidopsis [20], which may differentially impact on PIN polarization in columella cells. Accordingly, it would be interesting to investigate whether *LAZY/DRO* genes show indeed stage dependent regulation in LRs.

In conclusion, we substantiate our previous findings [7] and illustrate the preferential roles of PIN3 and PIN7 in early (stage I to II) and late (stage III to IV) gravity response of LRs, respectively.

## Materials and methods

### Plant material and growth conditions

*Arabidopsis thaliana* (L.) Heyhn. seedlings of Columbia ecotype (Col-0) were used in this study. The following lines have been previously described: DR5rev::GFP [21], PIN3::PIN3-GFP [22], PIN7::PIN7-GFP, PIN4::PIN4-GFP [11], pin3-5 pin7-1 [23], 35S::YUC8 [24] and 35S::YUC9 [24]. Seeds were sterilized with ethanol, plated on half Murashige and Skoog medium (1/2 MS salts (Duchefa), pH 5.9, 1% sucrose, and 0.8% agar), stratified at 4 °C for 2 days in the dark, and then transferred to a plant growth room (20-22°C, 16 h light/8 h dark, vertical orientation). The same medium and growth conditions were used for seedlings grown in 68x100 mm cylindrical vessels (Greiner), except for the use of a transparent agar (Gelrite, Duchefa) instead of traditional plant agar.

### Chemicals and treatments

Treatments with 1-Naphtaleneacetic acid (NAA) (Duchefa) were performed on 4-days-old seedlings (transferred to supplemented media). NAA was dissolved in DMSO before addition to MS+ medium. Confocal and fluorescence microscope imaging were performed 3 days after treatment. Scanning of plates for plateau length/PE angle measurements were performed 8-10 days after treatments. All experiments were performed at least three times.

### Microscopy

To detect PIN3-GFP, PIN4-GFP and PIN7-GFP signals in the tip of individual LRs, a binocular fluorescence microscope (Leica MZ FLIII) was used (GFP2 filters set: excitation filter 480/40 nm, barrier filter 510 nm) in combination with a DC 500 camera (Leica). To detect DR5rev::GFP and assess asymmetry of PIN7-GFP signal, confocal microscopy was performed using a Leica DM6000 CS, 347 TCS AOBS confocal laser scanning microscope, equipped with a HCX PL APO CS 348 20.0 0.70 IMM UV objective (excitation 488 nm, emission peak 509 nm). GFP signal in confocal images was analyzed and quantified using the software LAS AF 3.1 (Leica) and ImageJ (http://rsb.info.nih.gov/ij). All experiments were performed at least three times.

### Measurements

Plates with 12-14 DAG seedlings were scanned and the plateau length and plateau exit (PE) angle of individual LRs were measured using the line and angle tools of Image J, respectively. PE angles were measured using the head of the gravity vector as reference. GSA values at PE were sorted into 4 categories: 0˚-10˚, 11˚-20˚, 21˚-30˚, and 31˚-40˚. Percentages of incidence were calculated to generate graphs of PEA distribution. Values for ratio LR/plateau length were obtained by dividing the total length of the lateral root by the length of its plateau phase. Mean intensity of auxin signalling in the tip of LRs, as reported by the marker line DR5rev::GFP, was quantified using the polygon tool of ImageJ, circumscribing an area comprising the columella and lateral root cap cells, up to the height of the quiescent centre. For the assessment of PIN7-GFP asymmetry across the root tip, the mean intensity of the GFP signal was estimated for lateral plasma membrane domains of individual columella cells facing the upper and lower sections, or flanks, of the lateral root cap. Values deriving from the same section (LF and UF in Figure 3D, E) were averaged for each individual LR analyzed. “Polarization towards gravity” was defined when average mean intensity PIN7-GFP values for membrane domains in lower flank (LF) were bigger than average of mean intensity PIN7-GFP values for membrane domains in upper flank (UF). All experiments were performed at least three times.

### Statistics

The test of Kolmogorov-Smirnov (KS-test) (http://www.physics.csbsju.edu/stats/KS-test.n.plot_form.html) was used to evaluate statistical significance of differences observed between PE distributions. Student’s *t*-test was used for two-group comparisons. ANOVA, followed by Tukey‘s test, was used in the case of multiple comparisons to statistically evaluate differences between data sets. Sample size and usage of the tests are depicted in figure legends.

## Acknowledgements

We are grateful to J. Friml, S. Pollmann, S. Sabatini, and B. Scheres for sharing published material; Jit Thacker for help with preparing the manuscript; and the BOKU-VIBT Imaging Centre for access and expertise. This work was supported by the Vienna Research Group (VRG) program of the Vienna Science and Technology Fund (WWTF), the Austrian Science Fund (FWF) (Project: P29754), the European Research Council (ERC) (Starting Grant 639478-AuxinER) (to J.K-V.).

